# Yeast U6 snRNA made by RNA polymerase II is less stable but functional

**DOI:** 10.1101/2022.06.24.497417

**Authors:** Karli A. Lipinski, Jing Chi, Xin Chen, Aaron A. Hoskins, David A. Brow

**Author notes:** CORRESPONDING AUTHORS: Aaron A. Hoskins,; David A. Brow.

## Abstract

U6 small nuclear (sn)RNA is the shortest and most conserved snRNA in the spliceosome and forms a substantial portion of its active site. Unlike the other four spliceosomal snRNAs, which are synthesized by RNA polymerase (RNAP) II, U6 is made by RNAP III. To determine if some aspect of U6 function is incompatible with synthesis by RNAP II, we created a U6 snRNA gene with RNAP II promoter and terminator sequences. This “U6-II” gene is functional as the sole source of U6 snRNA in yeast, but its transcript is much less stable than U6 snRNA made by RNAP III. Addition of the U4 snRNA Sm protein binding site to U6-II increased its stability and led to formation of U6-II•Sm complexes. We conclude that synthesis of U6 snRNA by RNAP III is not required for its function and that U6 snRNPs containing the Sm complex can form *in vivo*. The ability to synthesize U6 snRNA with RNAP II relaxes sequence restraints imposed by intragenic RNAP III promoter and terminator elements and allows facile control of U6 levels via regulators of RNAP II transcription.

## INTRODUCTION

The five small nuclear RNAs U1, U2, U4, U5, and U6 are essential components of the eukaryotic spliceosome, necessary both for precise identification of the intron-exon boundaries and splicing catalysis (Guthrie and Patterson 1988). Of the five spliceosomal snRNAs, four are synthesized by RNA polymerase (RNAP) II. Only U6, the smallest of the spliceosomal RNAs, is synthesized by RNAP III (Didychuk et al. 2018; Dergai and Hernandez 2019). Indeed, it appears that RNAP II is actively excluded from the vicinity of the *Saccharomyces cerevisiae* yeast U6 snRNA gene, *SNR6* (Steinmetz et al. 2006). It is not known why U6 is made by RNAP III, but it could be because some characteristic of RNAP II synthesis is inconsistent with U6 snRNA function. For example, RNAP III typically starts transcription at a single position, defined primarily by spacing from an intragenic “A block” promoter element (Gerlach et al. 1995), whereas RNAP II often starts at multiple positions defined by a less stringent initiator element (Kuehner and Brow 2006). Also, RNAP II transcripts receive a reverse 7-methylguanosine “cap” on their 5′ triphosphate, whereas RNAP III transcripts generally do not. Human U6 snRNA instead receives a γ-monomethylphosphate cap on its 5′ triphosphate, which may be present in yeast and other organisms as well (Didychuk et al. 2018). However, yeast U6 molecules made by RNAP III but bearing altered 5′ ends and methylguanosine caps are functional in vivo (Kwan et al. 2000).

Another possible reason that U6 snRNA is made by RNAP III is to promote assembly of the Lsm2-8 protein ring on its 3′ tail (Mayes et al. 1999; Karaduman et al. 2006). All transcripts made by RNAP III terminate with an oligo(U) tail of four or more residues (Arimbasseri and Maraia 2015). On yeast U6 snRNA the oligo(U) tail, which initially has a terminal 2’,3’ *cis* diol, is bound by the La-homologous protein Lhp1 (Pannone et al. 1998; Didychuk et al. 2017). Subsequently, the terminal nucleoside of the oligo(U) tail is removed by the 3’-5’ exonuclease Usb1 leaving a 3’ phosphate group on the new terminal nucleotide. The trimmed tail is then bound by the Lsm2-8 heteroheptameric protein ring (Mayes et al. 1999; Didychuk et al. 2017; Montemayor et al. 2018). Retention of mature U6 in the nucleus is dependent on binding of Lsm2-8 (Spiller et al. 2007a,b). In contrast, the 3′ tails of the spliceosomal snRNAs made by RNAP II instead bind to a paralogous, seven-protein Sm ring, which assembles on an AUUUUUG sequence internal to the 3′ terminus (Jones and Guthrie 1990; Salgado-Garrido et al. 1999).

To determine if synthesis of U6 snRNA by RNAP III is required for its function, we replaced the upstream and downstream sequences of *SNR6* with RNAP II control elements. Here we report that U6 snRNA synthesized by RNAP II (U6-II) is functional *in vivo* but is highly unstable. U6 snRNA made from a glucose-repressible variant of the U4 snRNA (*SNR14*) promoter has a half-life of less than 15 minutes, as compared to an estimated half-life of more than 6 hours for U6 made by RNAP III. We hypothesized that the instability of U6-II is due to inefficient recruitment of the Lsm protein complex. Consistent with this hypothesis, addition of the Sm binding site of U4 snRNA to the 3’ tail of U6-II (U6-II-Sm) strongly stabilizes it, increasing the half-life to about 2 hours. Binding of the Sm ring to U6-II-Sm was confirmed by co-immunoprecipitation of free U6 RNA with the Sm protein Smd1. We conclude that biosynthesis by RNAP II is compatible with yeast U6 snRNA function. The ability to control U6 snRNA synthesis with regulated RNAP II promoters expands the tools available to probe U6 biogenesis and function.

## RESULTS

### Functional U6 snRNA can be made by RNAP II

*SNR6* has an unusual RNAP III promoter structure, with upstream, intragenic, and downstream elements (Eschenlauer et al. 1993; **Fig. 1A**). To convert U6 snRNA into an RNAP II transcript, which we call U6-II, we replaced the upstream and downstream DNA flanking the U6 snRNA-coding region with the promoter and terminator from the RNAP II-transcribed U4 snRNA gene, *SNR14* (Kuehner and Brow 2006; Steinmetz et al. 2001). Our initial construct contained *SNR14* sequence from -224 to +701 relative to the transcription start site (TSS, at position +1), but with the 160-nucleotide U4 snRNA coding region replaced with the 112-nucleotide U6 snRNA coding region. Note that the intragenic A block promoter element, while present in this construct, is not able to direct RNAP III transcription of the U6 gene in the absence of the downstream B block element (Brow and Guthrie 1990; **Fig. 1A**). This construct, named *SNR14-6-14*, did not support viability in the absence of *SNR6* unless present on a high-copy plasmid.

**Figure 1.**
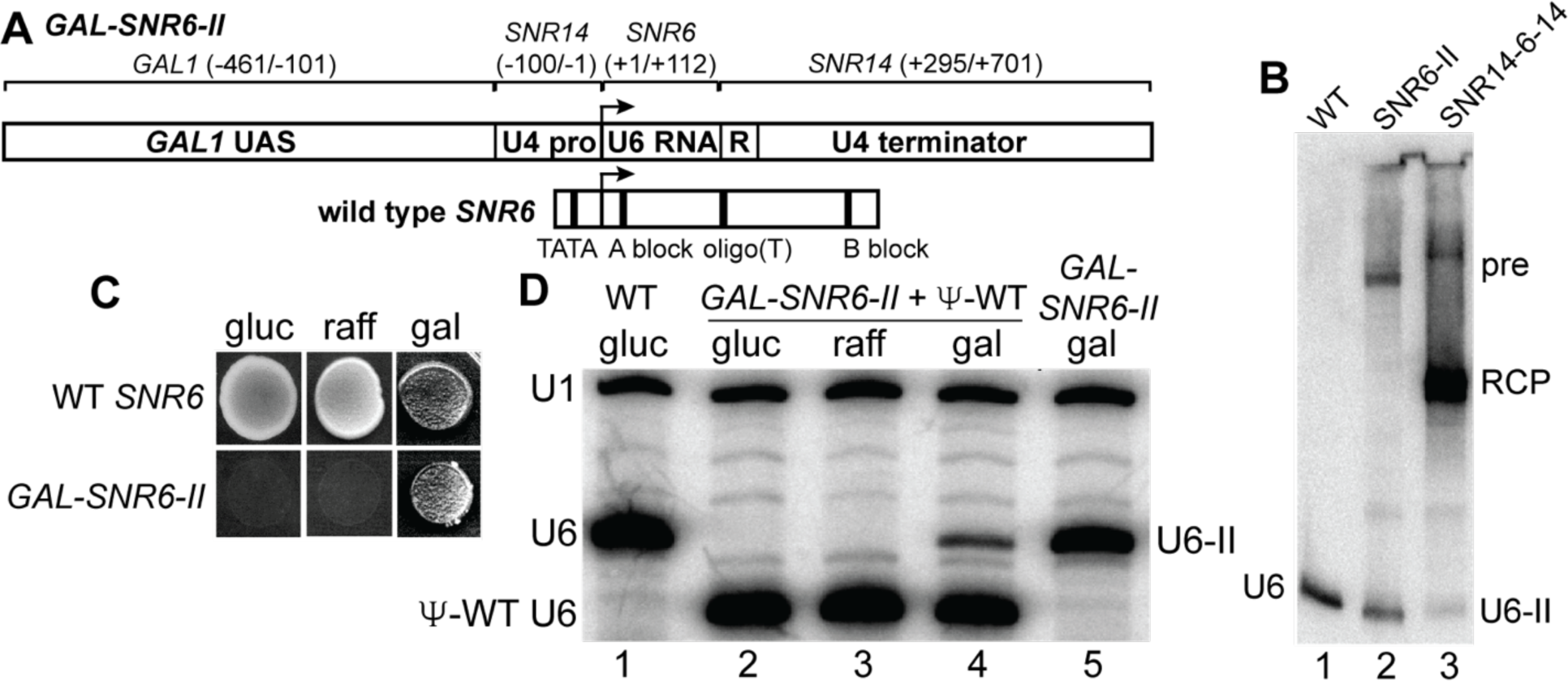
Expression of functional yeast U6 snRNA by RNAP II. **A)** Schematic of the *GAL-SNR6-II* allele. The *GAL1* upstream activating sequence (UAS) was fused to the U4 snRNA gene (*SNR14*) TATA box and proximal promoter (U4 pro), followed by the coding sequence of the U6 snRNA gene (*SNR6*). The terminator includes the Rnt1 site (R) and downstream sequences of *SNR14*. The wild type *SNR6* gene is shown below for comparison, with the RNAP III promoter elements and oligo(T) terminator indicated by thick vertical lines. **B)** Northern blot of total cellular RNA from an *SNR6* disruption strain (MWK027) containing a high-copy plasmid (pRS424) bearing the WT *SNR6*, *SNR6-II*, or *SNR14-6-14* allele of the U6 snRNA gene, using a probe complementary to the 3’ end of U6 snRNA. pre: presumptive primary transcript; RCP: presumptive Rnt1 cleavage product; U6-II: U6 snRNA synthesized by RNAP II. **C)** Growth of an *SNR6* disruption strain (MWK027) containing the indicated *SNR6* allele on a low-copy plasmid (pRS314) on medium containing glucose (gluc), raffinose (raff), or galactose (gal). **D)** Primer extension analysis of U1 and U6 snRNAs in a U6 disruption strain (MWK027) containing the indicated *SNR6* allele(s) on low-copy plasmid(s) and grown in medium containing the indicated sugars. A shortened “pseudo-WT” (Ψ-WT) U6 allele is present in lanes 2-4 to provide U6 function in conditions where *GAL-SNR6-II* is repressed. Note that the level of U6-II is increased in the absence of Ψ-WT U6. U1 RNA serves as a normalization control.

Northern blot analysis of RNA produced *in vivo* from the high-copy *SNR14-6-14* construct indicated that most of the U6 transcript was 3’-extended, with the major species 250-300 nucleotides in length (**Fig. 1B**, lane 3, “RCP”). This length corresponds to the known position of a Rnt1 cleavage site important for the 3’-processing of the pre-U4 snRNA (Allmang et al. 1999). Normally, the Rnt1 cleavage product of pre-U4 snRNA does not accumulate significantly, due to rapid 3’-trimming by the exosome. However, it is possible that 3’-trimming is inhibited in the context of the U6 snRNA coding sequence. To increase the yield of U6-II with the proper 3’ end, we deleted the first 135 nucleotides of *SNR14* sequence downstream of the U6 snRNA coding region, which places the upstream Rnt1 cleavage site at position +112 of the U6 snRNA coding region (**Figs. 1A** and **S1**). We named this construct *SNR6-II*. When expressed from a high copy-number plasmid in yeast cells, the *SNR6-II* allele produced U6 snRNA of the correct size (U6-II), as well as a 3’-extended product corresponding in length to uncleaved primary transcript (**Fig. 1B**, lane 2).

To generate a regulated version of the *SNR6-II* construct with a potentially higher expression level, we deleted DNA upstream of the *SNR14* TATA box (upstream of +100) and replaced it with the 461 base pairs of DNA immediately upstream of the *GAL1* TATA box (**Fig. 1A**). This substitution put the *SNR6-II* construct under control of the galactose-inducible activator Gal4 (Johnston and Davis 1984). We found that the *GAL-SNR6-II* construct on a low copy-number plasmid rescues deletion of the chromosomal U6 gene, but only on growth medium containing galactose and not on medium containing glucose or raffinose, which are expected to repress or not induce Gal4-regulated genes, respectively (**Fig. 1C**).

To confirm expression or repression of the Gal4-regulated U6 gene, we generated a merodiploid strain containing two different *SNR6* alleles: *GAL-SNR6-II* and a “pseudo-wild type” (Ψ-WT) U6 gene containing a shortened 5′ stem-loop and under control of the native RNAP III promoter (Madhani et al. 1990). Ψ-WT U6 maintains yeast viability in the presence of raffinose or galactose and allows quantification of the levels of U6-II as the two RNAs are of different lengths. Primer extension analysis confirmed that no detectable U6-II was expressed from the *GAL-SNR6-II* construct in the presence of glucose or raffinose (**Fig. 1D**, lanes 2 and 3) and explains why yeast failed to grow under those conditions without Ψ-WT U6.

Primer extension analysis also showed that the low copy-number *GAL-SNR6-II* construct produces about one-third the WT amount of U6 snRNA in the presence of medium containing galactose (**Fig. 1D**, *cf*. lanes 1 and 5). We showed previously that a similar amount of RNAP III-made U6 snRNA is sufficient for normal growth (Kaiser and Brow 1995). It is interesting that the steady-state level of U6-II (relative to U1 snRNA) is diminished in the presence of Ψ-WT U6 (**Fig. 1D**, *cf.* lanes 4 and 5). This observation suggests that either the two U6 genes are competing for transcription factors, which should not be the case since they are presumably transcribed by different RNAPs, or that the RNAs are competing for binding proteins that stabilize them. The latter is more likely since both should bind Prp24 and the Lsm2-8 ring (Montemayor et al. 2018). If U6-II is out-competed by Ψ-WT U6 for these proteins, then it may be destabilized and degraded leading to the lower steady-state level. Weaker binding of the Lsm2-8 ring to U6-II could be due to lack of 3’-end processing by Usb1 to produce a terminal 3’-phosphate or shortening of the U6 Lsm binding site from four to three or fewer uridines by Usb1. In either case the binding affinity of U6-II for Lsm2-8 would be expected to be reduced (Didychuk et al. 2017) and result in a competitive disadvantage for protein binding by U6-II. Importantly, the combined observations show that a functional U6 snRNA can be made by RNAP II using a galactose-inducible promoter. Thus, transcription by RNAP III is not required for U6 function.

### U6 RNA made by RNAP II is methylguanosine-capped

Human U6 snRNA is *O*-methylated on the γ-phosphate group of its initiating GTP residue (Singh and Reddy 1989). It is not known if yeast U6 snRNA receives the same modification, but it is known that disruption of the recognition element for the human U6 capping enzyme in yeast U6 results in the acquisition of a RNAP II-like monomethylguanosine (MMG) 5’-cap *in vivo*, even when the U6 is produced by RNAP III (Kwan et al. 2000). This MMG cap is subsequently hypermethylated to a trimethylguanosine (TMG) cap. Since methylguanosine capping of RNAP II transcripts normally occurs cotranscriptionally, we would expect some fraction of U6-II to receive a MMG cap, even though it has an intact determinant for a potential γ-*O*-methylphosphate cap. We assessed methylguanosine-capping of U6-II by antibody gel shift with the monoclonal antibody (mAb) H20 that recognizes both the TMG and, to a lesser extent, MMG cap (Bochnig et al. 1987). U4 snRNA acts as an internal control, and its mobility is efficiently shifted by binding of H20 mAb (**Fig. 2**, *cf.* lanes 1, 2, 5, and 6). RNAP III-made U6 exhibits no binding to H20 mAb, as expected (*cf.* lanes 3 and 4), but RNAP II-made U6 is partially shifted by H20 mAb (*cf.* lanes 7-10). The incomplete shifting of U6-II could be due to competing γ-*O*-methylphosphate capping, incomplete hypermethylation of MMG to TMG cap, or occlusion of mAb binding by the U6 5’ stem-loop (Kwan et al. 2000).

**Figure 2.**
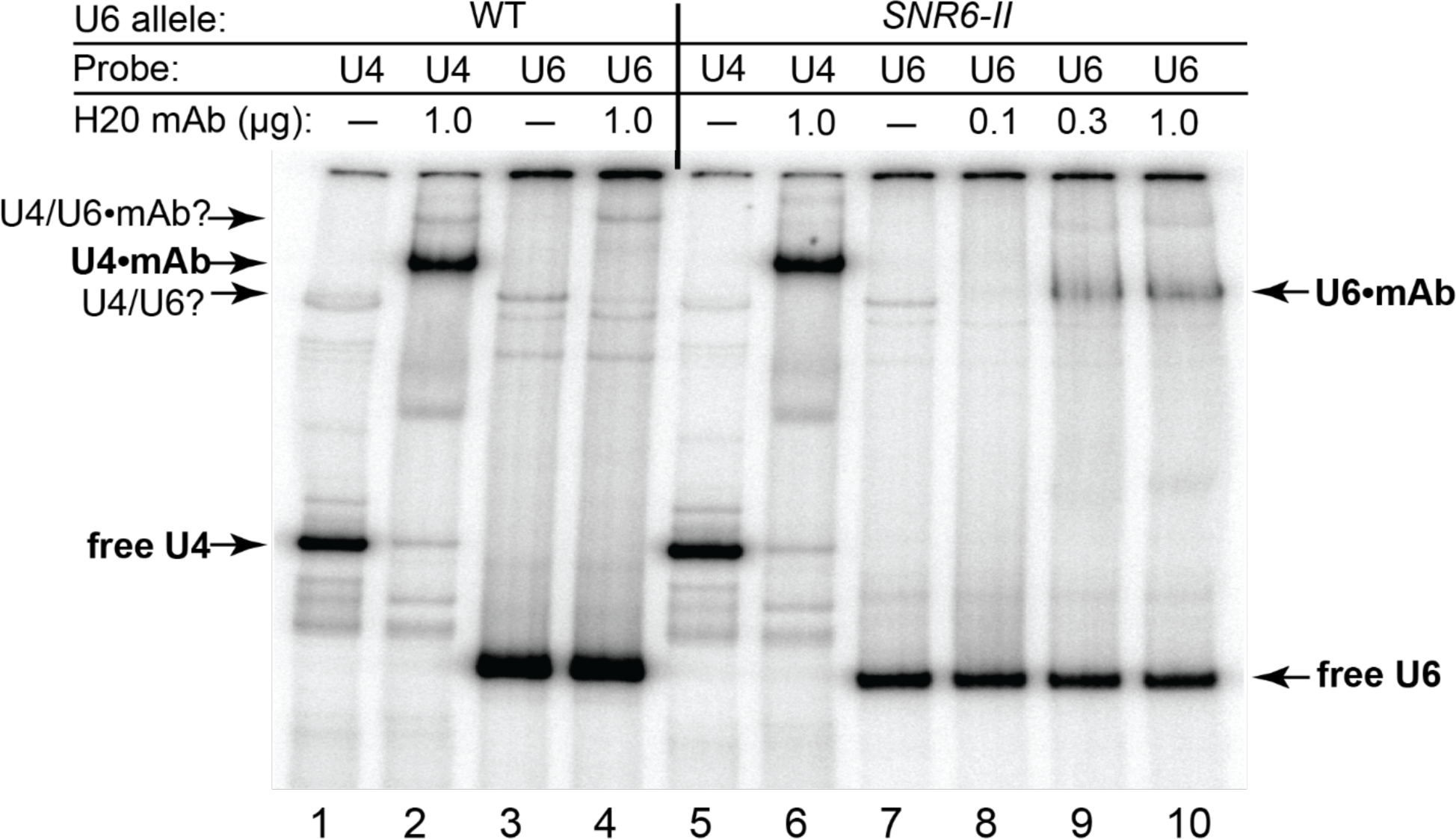
Evidence for methylguanosine-capping of U6 snRNA made by RNAP II. Total RNA from MWK027 with pRS314-*SNR6* (WT) or pRS314-*GAL-SNR6-II* (*SNR6-II*) was heated for 1 min at 90°C,s then hybridized to ^32^P-labeled oligonucleotide probes specific for U4 or U6 snRNA. The RNA was then incubated with the indicated amount of mAb H20, which binds the TMG or MMG caps, prior to electrophoresis on a non-denaturing gel. The positions of free U4 and U6 snRNAs, and mAb-shifted snRNAs are indicated. A minor band consistent with base-paired U4/U6 is also seen in both samples.

### U6 snRNA made by RNAP II is highly unstable

Creation of a U6 gene under control of the *GAL1* UAS made it possible to shut off U6 snRNA synthesis by switching the carbon source from galactose to glucose, which allowed us to measure the *in vivo* half-life of U6-II. When transferred to glucose medium, yeast cells bearing the *GAL-SNR6-II* construct stopped growing in about 10 h (**Fig. 3A**), more rapidly than yeast cells bearing a Gal4-regulated U1, U2, or U5 RNA gene (Patterson and Guthrie 1987; Seraphin and Rosbash 1989). Analysis of U6 snRNA from the *GAL-SNR6-II* strain revealed a half-life of 12 min for U6-II (**Figs. 3B, C**). It has not been possible to measure a precise half-life of RNAP III-made U6 RNA, but studies using the *lac* repressor to block RNAP III transcription (Luukkonen and Seraphin 1998) or a temperature-sensitive mutation in a RNAP III subunit (Kwan et al. 2000) suggest that RNAP III-made U6 has a half-life of greater than 6 h. Thus, U6-II is at least 30-fold less stable than U6 snRNA made by RNAP III.

**Figure 3.**
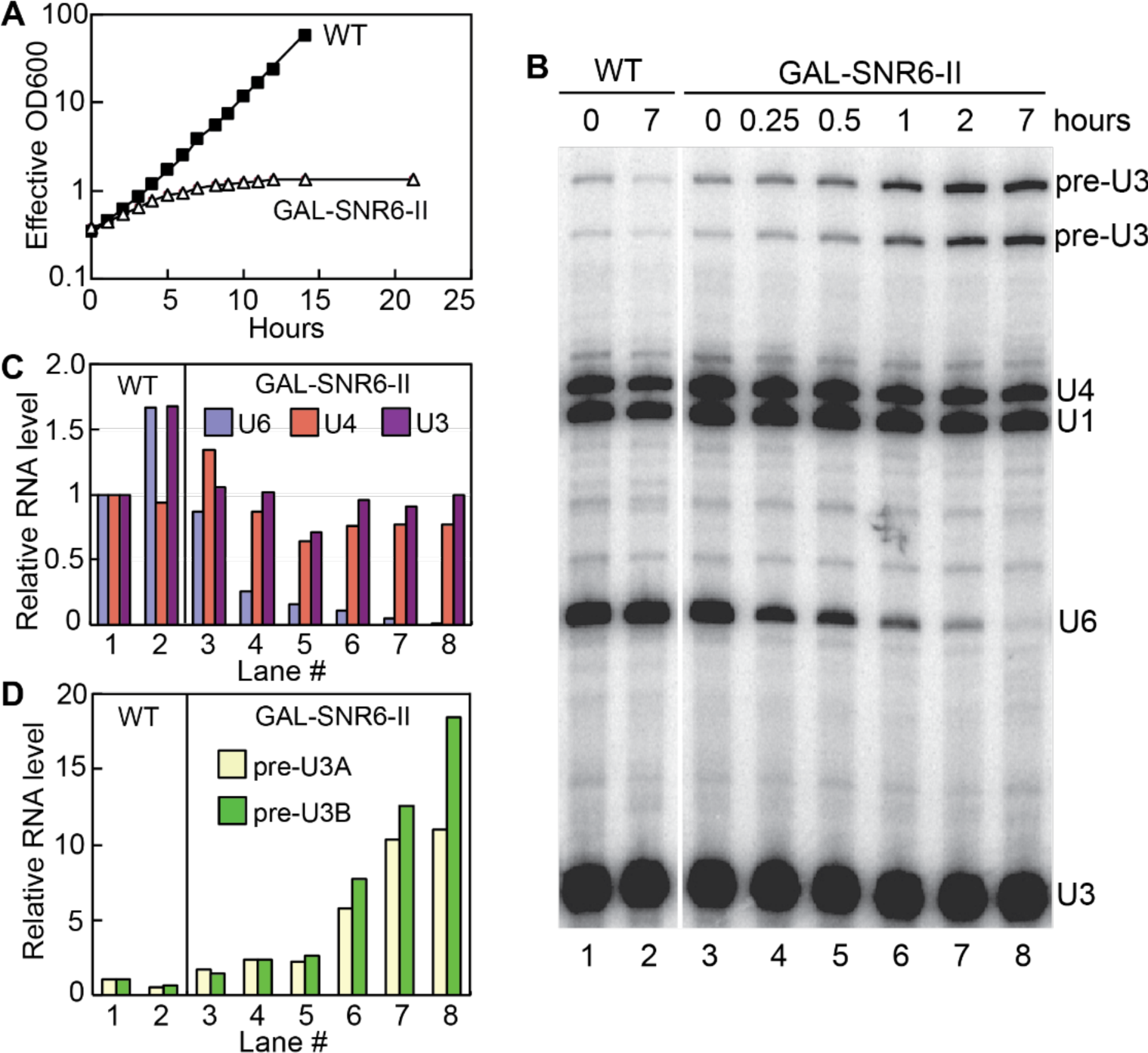
U6 snRNA made by RNAP II is unstable in vivo. **A)** Growth curve of an *SNR6* disruption strain (MWK027) containing pRS314-*SNR6* (WT) or pRS314-*GAL-SNR6-II* after replacement of YEP-galactose medium with YEP-glucose at time 0 to repress transcription of the *GAL-SNR6-II* allele. Both strains were diluted to keep the optical density at 600 nm (OD600) less than 1. The “effective OD600” value is corrected for this dilution. **B)** Total RNA extracted from cells collected at the indicated times was subjected to primer extension to detect U1, U4, and U6 snRNAs as well as U3 snoRNA, which is transcribed from two loci (A and B) as an intron-containing precursor (pre-U3). **C)** Quantification of U6, U4, and mature U3 RNA present in the indicated lanes of the gel from part B relative to the WT strain immediately before shift from galactose to glucose (Lane 1). Values are normalized to U1 snRNA in each sample to control for variable RNA recovery. **D)** Quantification of pre-U3 snoRNA present in the indicated lanes of the gel from part B relative to the wildtype strain immediately before shift from galactose to glucose (Lane 1). Values are normalized as in Panel C.

We monitored splicing as a function of U6 snRNA abundance by measuring the level of pre-U3 snoRNA, which is made from two genes (*SNR17A* and *SNR17B*) that have spliceosomal introns of different lengths. In the presence of galactose, the steady state level of pre-U3 RNA in the *GAL-SNR6-II* strain is essentially the same as in a WT strain (**Figs. 3B, D**). However, as soon as an hour after shift to glucose, an increase in pre-U3 is observed in the *GAL-SNR6-II* strain. (Because mature U3 snoRNA is very stable, little change in its level is observed over the time course.) Thus, the rapid kinetics of sglucose repression and U6-II turnover result in quick disablement of the spliceosome in response to the change in medium.

### U6-II is stabilized by inclusion of an Sm protein-binding sequence

The spliceosomal RNAs that are normally synthesized by RNAP II bind the heteroheptameric Sm protein ring, which contributes to the stability of these RNAs (Jones and Guthrie 1990). In contrast, U6 snRNA is bound and stabilized by the paralogous Like-Sm (Lsm) heteroheptamer (Mayes et al. 1999). Binding of the Lsm proteins to U6 snRNA may be coupled to RNAP III transcription via the La-homologous protein, Lhp1, which binds the oligo(U) tail of RNAP III transcripts (Pannone et al. 1998, 2001). In addition, the affinity of Lsm2-8 for U6 is determined by post-transcriptional processing of the 3’ end of U6 by Usb1, as mentioned previously. Therefore, it is not clear if U6-II would efficiently bind and be stabilized by the Lsm proteins (but see below).

To test if binding of Sm proteins could stabilize U6-II and is compatible with U6 snRNA function, we replaced 11 nucleotides at the 3’ end of U6 with 14 nucleotides from the 3’ end of U4, creating the *GAL-SNR6-II-Sm* allele (**Fig. 4A**). This substitution creates a consensus Sm binding site (A2U5G2) beginning at position 98 of the U6 snRNA but retains a four-uridine Lsm binding site at the 3’ end. A strain bearing the *GAL-SNR6-II-Sm* allele on a low copy plasmid was viable, indicating that U6-II-Sm is functional. RNA analysis showed that U6-II-Sm is much more stable than U6-II, with a half-life of about two hours (**Fig. 4B**, **C**). Two hours after transfer from galactose to glucose medium about five times as much U6-II-Sm (relative to U1 snRNA) is present than U6-II, yet the U6-II-Sm strain accumulated roughly twice as much unspliced pre-U3 snoRNA, suggesting that U6-II-Sm may not be as functional as U6-II. Thus, creation of an Sm-binding site in the 3’ tail of U6-II increases its stability but may decrease its function.

**Figure 4.**
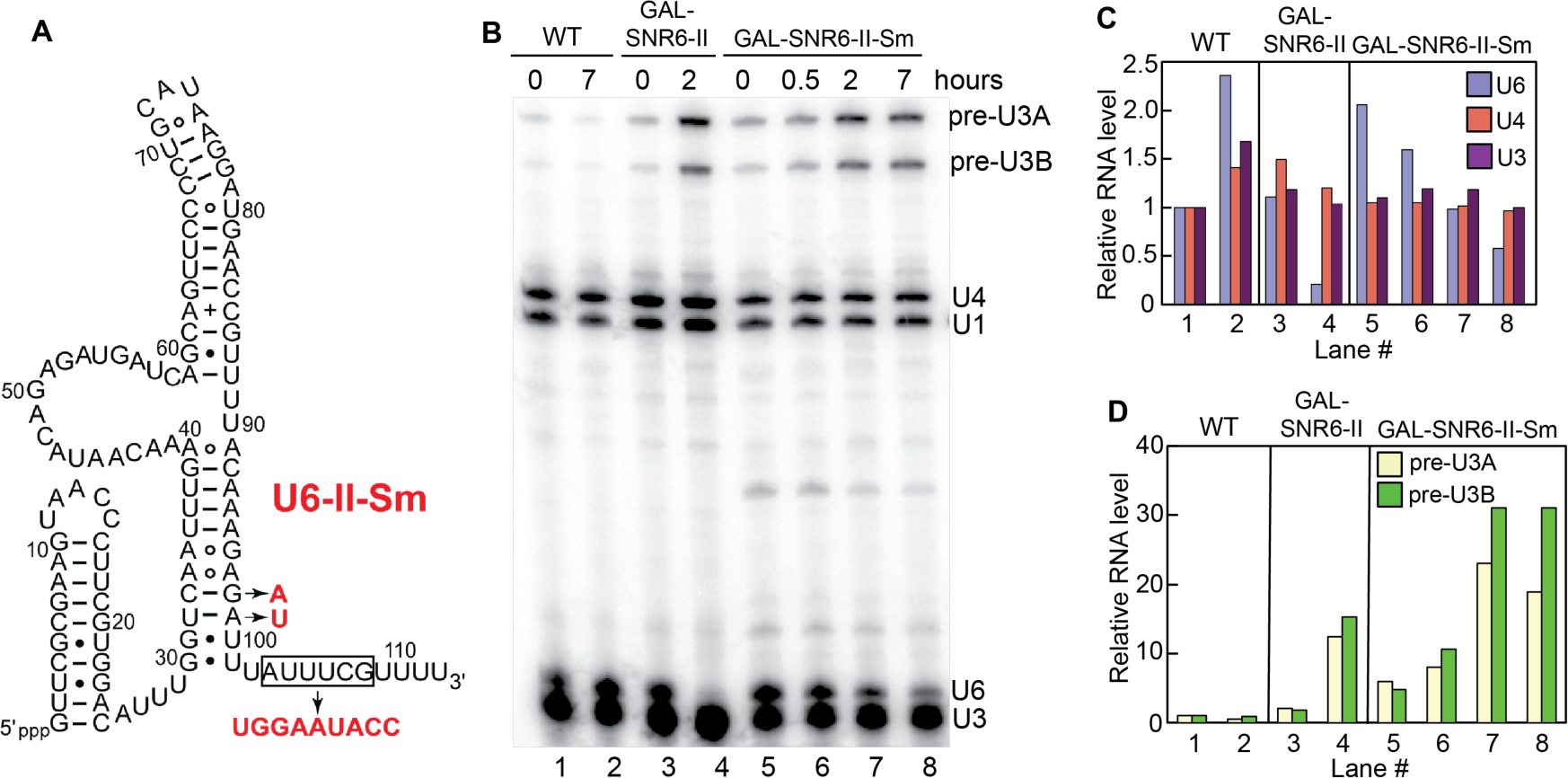
Creation of a consensus Sm-binding site in U6-II increases its stability. **A)** Substitutions (in red) introduced in U6-II to replace its 3’ end with that of U4 snRNA, including the Sm-binding site (A2U5G2). Replacement of six nucleotides of U6 with nine of U4 elongate U6-II-Sm by three nucleotides. **B)** Total RNA extracted from cells with WT U6, U6-II or U6-II-Sm was collected at the indicated times and subjected to primer extension to detect U1, U4, and U6 snRNAs as well as unspliced and spliced U3 snoRNA. (Note that a different U6 primer, U6B, was used than in Figure 3.) **C)** Quantification of U6, U4, and mature U3 RNA present in the indicated lanes of the gel from part B relative to the wildtype strain immediately before shift from galactose to glucose (Lane 1). **D)** Quantification of pre-U3 snoRNA present in the indicated lanes of the gel from part B relative to the wildtype strain immediately before shift from galactose to glucose (Lane 1). Values are normalized to U1 snRNA in each sample to control for variable RNA recovery.

### Free U4 snRNP accumulates in the presence of U6-II

To examine how U6-II impacts the distribution of snRNAs among splicing complexes, extracts made from yeast expressing either WT U6, U6-II, or U6-II-Sm were sedimented through a glycerol gradient to resolve different snRNPs and spliceosomes. Primer extension of the RNA extracted from each gradient fraction was then used to identify the snRNAs present (**Fig. S2**). It should be noted that the haploid yeast strain used in these experiments also contained a deletion of the *SNR14* gene with U4 snRNA being provided from a low-copy plasmid, so U4 gene dosage could potentially increase under selective pressure.

U6 assembly into spliceosomes normally requires base pairing with U4 to form a di-snRNP particle, which then binds the U5 snRNP to form the U4/U6.U5 tri-snRNP.

Therefore, the U4 to U6 ratio is expected to be equal to one in fractions containing only tri-snRNP. Fractions at the bottom of the gradient (24 and 25) have U4/U6 ratios of less than one relative to tri-snRNP fractions, consistent with activated spliceosomes that have lost U4 snRNP (**Fig. 5**).

**Figure 5.**
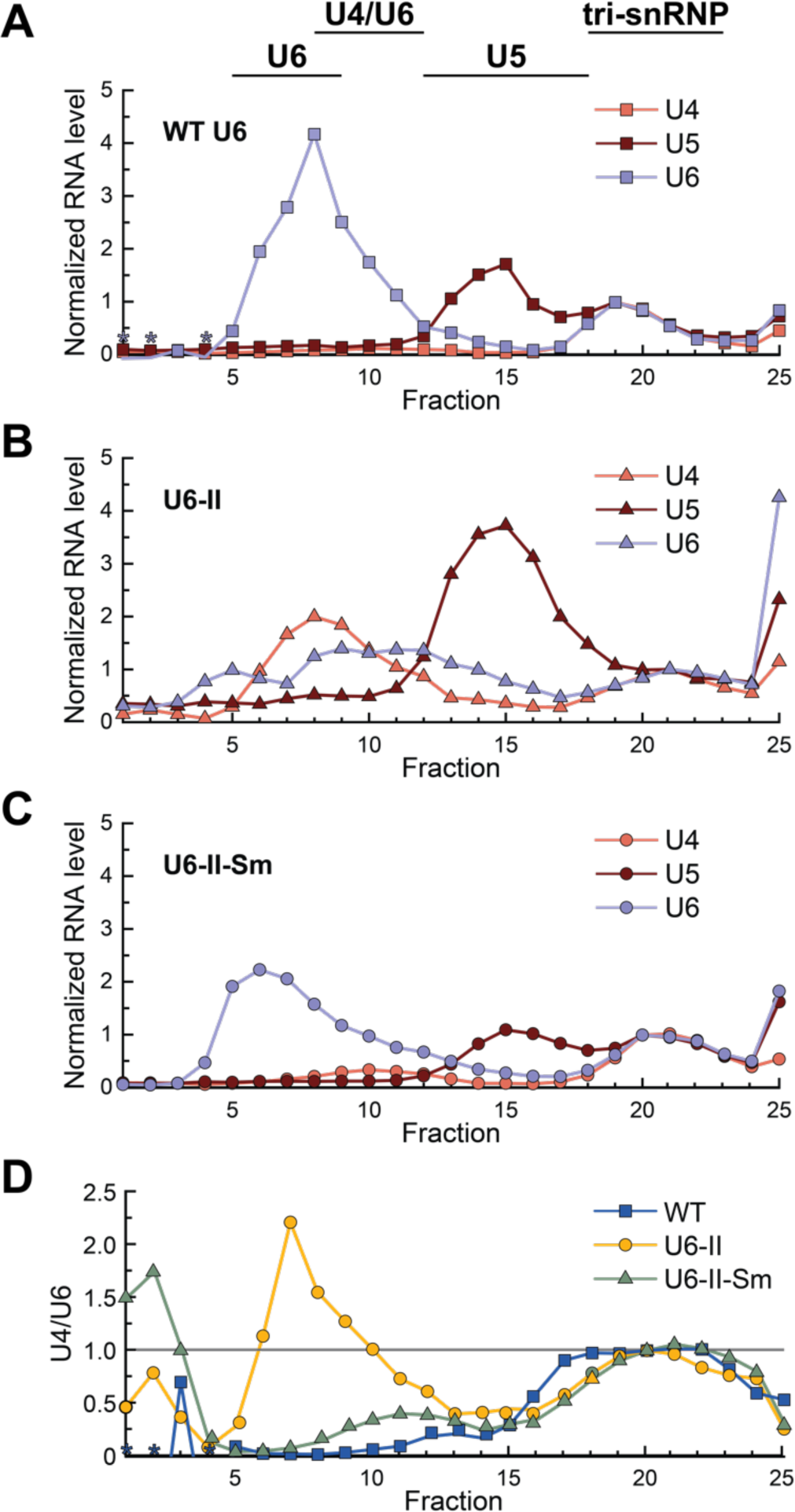
Glycerol gradient fractionation reveals altered snRNP distributions in U6-II extracts. Yeast cell extracts were sedimented through a 10-30% glycerol gradient, fractionated, and snRNAs analyzed by primer extension. To normalize the data, the U4, U5 and U6 primer extension products in peak tri-snRNP-containing fractions for each gradient (20 or 21) were set to values of “1” since these RNAs are expected to be equimolar in tri-snRNPs. Asterisks denote fractions for which no detectable U4 RNA above background was detected. **A-C)** Relative abundances of the U4 (red), U5 (brown), and U6 (blue) RNAs in each glycerol gradient fraction obtained using WT (A), U6-II (B), or U6-II-Sm (C) yeast extract. **D)** The normalized U4/U6 ratio is plotted for each fraction from gradients of extracts prepared from WT (blue squares), U6-II (yellow circles), or U6-II-Sm (green triangles) strains.

In cell extract with WT U6, there is a large excess of U6 over U4 and the U4 to U6 ratio is very low in fractions not containing tri-snRNP (**Figs. 5A, 5D, and S2**). These results are consistent with previous studies of snRNP distribution in WT yeast extracts (Bordonné et al. 1990). In contrast, extract with U6-II exhibited increased levels of free U4 snRNP (**Figs. 5B and 5D**, fractions 6-10) and free U5 snRNP (**Fig. 5B**, fractions 13-18), indicating inefficient formation of U4/U6 di-snRNP and U4/U6.U5 tri-snRNP. U6-II extract also showed a large amount of U6 sedimenting at the bottom of the gradient (**Fig. 5B**, fraction 25). When the U6-II-Sm extract was analyzed, the snRNA distribution in gradient fractions much more closely resembled the distribution seen with WT U6 (*cf*. **Figs. 5A** and **5C**).

Accumulation of U4 snRNA that is not base paired to U6-II was also observed by solution hybridization of radiolabeled oligonucleotides to cold-extracted cellular RNAs followed by nondenaturing gel electrophoresis (**Fig. S3**). RNA prepared from the U6-II strain contained a higher percentage of U4 not paired to U6 than either the U6-II-Sm or WT U6 extracts (**Fig. S3B and S3F**). The total amount of U6 in the U6-II strain is also less than the amounts present in either WT U6 or U6-II-Sm (**Fig. S3A, S3C**). Excess free U4 snRNA was also observed when RNAP III-made U6 levels were reduced by extragenic promoter mutations (Kaiser and Brow 1995), so reduced U6-II stoichiometry may be the main reason for accumulation of free U4. However, we cannot rule out that U6-II assembles less efficiently into U4/U6 di-snRNP, for example, due to decreased Prp24 or Lsm2-8 binding (Didychuk et al. 2015).

### U6-II and U6-II-Sm are associated with Lsm proteins

It is possible that synthesis of U6 by RNAP II, with or without inclusion of an Sm protein binding site, could reduce or even eliminate Lsm2-8 protein binding. To test if this occurs, we first looked for genetic interactions between U6-II-Sm and Lsm proteins. Single-gene deletions of *lsm5* and *lsm8* are not lethal if U6 snRNA is over-expressed, consistent with the primary function of the Lsm2-8 proteins in yeast being stabilization of U6 snRNA (Pannone et al. 2001; Roth et al. 2018). We predicted that if Lsm5 or Lsm8 association with U6-II-Sm is not essential, then U6-II-Sm yeast cells should be viable in *lsm511* or *lsm8Δ* strains after selection for loss of complementing *URA3*-marked plasmids. Instead, the *lsm5* and *lsm8* deletions were lethal in cells expressing U6-II-Sm (**Fig. S4**). This result suggests that the Lsm2-8 ring associates with U6-II-Sm snRNA. It should be noted that *lsm5* and *lsm8* deletions were viable in the presence of WT U6 (**Fig. S4**), likely due to amplification of the low-copy plasmid containing the U6 gene (Pannone et al. 2001; Roth et al. 2018). The stronger requirement for Lsm5 and Lsm8 observed with U6-II-Sm may have to do with its decreased function, noted above.

To provide evidence for a physical interaction between the Lsm2-8 ring and U6-II or U6-II-Sm, we created yeast strains with an integrated, C-terminal 2xV5 epitope tag on the Lsm8 protein and used anti-V5 antibody to immunoprecipitate (IP) Lsm8-associated RNAs. In WT U6-expressing cells, the U4, U5, and U6 snRNAs but not the U1 and U2 snRNAs co-IP’ed with Lsm8-2xV5 (**Fig. 6**, lane 6). This result is consistent with Lsm2-8-bound U6 snRNA being a component of the U6 snRNP, U4/U6 di-snRNP, and the U4/U6.U5 tri-snRNP, while the Lsm ring is absent from activated or catalytic spliceosomes (Chan and Cheng 2005). U6-II and U6-II-Sm snRNAs were also co-IP’ed with Lsm8-2xV5 (**Fig. 6**, lanes 7 and 8). In both cases, U4 and U5 but not U1 and U2 RNAs were also present, albeit in lower amounts. These data support binding of U6-II and U6-II-Sm by Lsm8, likely as part of the Lsm2-8 ring. Thus, it appears that U6 synthesis by RNAP III is not required for Lsm2-8 protein binding and that the inclusion of an Sm-binding site into U6-II-Sm does not prevent Lsm ring binding.

**Figure 6.**
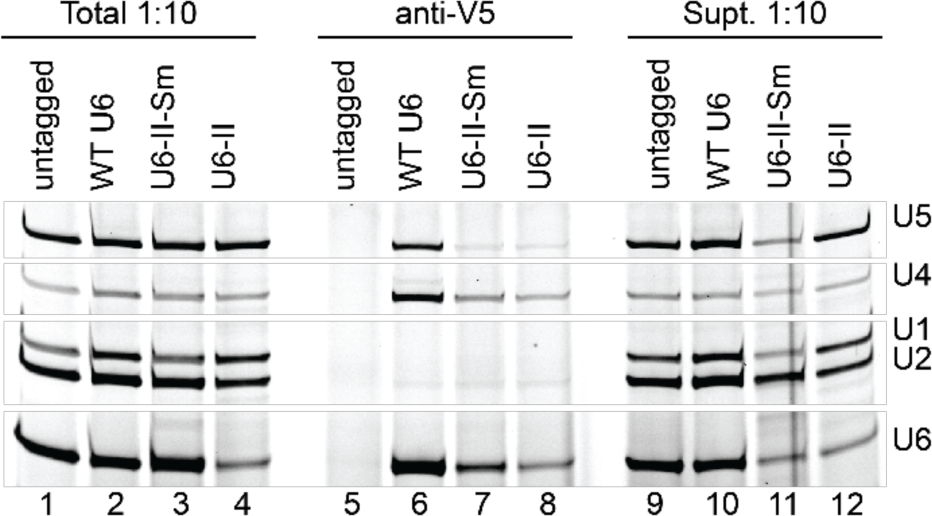
U6-II and U6-II-Sm co-IP with the Lsm8 protein. Extracts from Lsm8-2xV5 strains or a control lacking the 2xV5 epitope tag were immunoprecipitated using anti-V5 antibodies. RNAs from the immunoprecipitate, supernatant, or total extract were extracted using phenol, precipitated, and analyzed by primer extension.

### U6-II-Sm binds Sm proteins

The stability conferred to U6-II transcripts by the addition of an Sm binding site suggests that U6-II-Sm is bound by Sm proteins. To investigate whether the Sm ring binds U6-II-Sm, we created yeast strains with an integrated C-terminal 3xFLAG epitope tag on the Smd1 protein and used anti-FLAG antibody to IP Smd1-associated RNAs. Pulldown of WT U6 with Smd1 is possible due to its formation of complexes containing U1, U2, U4, and/or U5 with bound Sm rings. Not surprisingly, all five snRNAs co-IP’ed with Smd1 in all three strains (**Fig. 7A**). We quantified the ratio of U6 to U4 in the WT U6 sample and set the value equal to 1, since the RNAs should be stoichiometric in the U4/U6 and U4/U6.U5 snRNPs, with little free U4 snRNP present (**Fig. 5A**). In contrast, the U6 to U4 ratio in the U6-II-Sm IP (normalized to WT U6) was 1.5, indicating that a third of the U6-II-Sm in the IP may originate from free U6 snRNP that contains Smd1 (**Fig. 7B**). Notably, the same calculation for U6-II IP gave a U6/U4 ratio of ∼0.8, consistent with the presence of excess free U4 snRNP in this extract (**Fig. 5B**).

**Figure 7.**
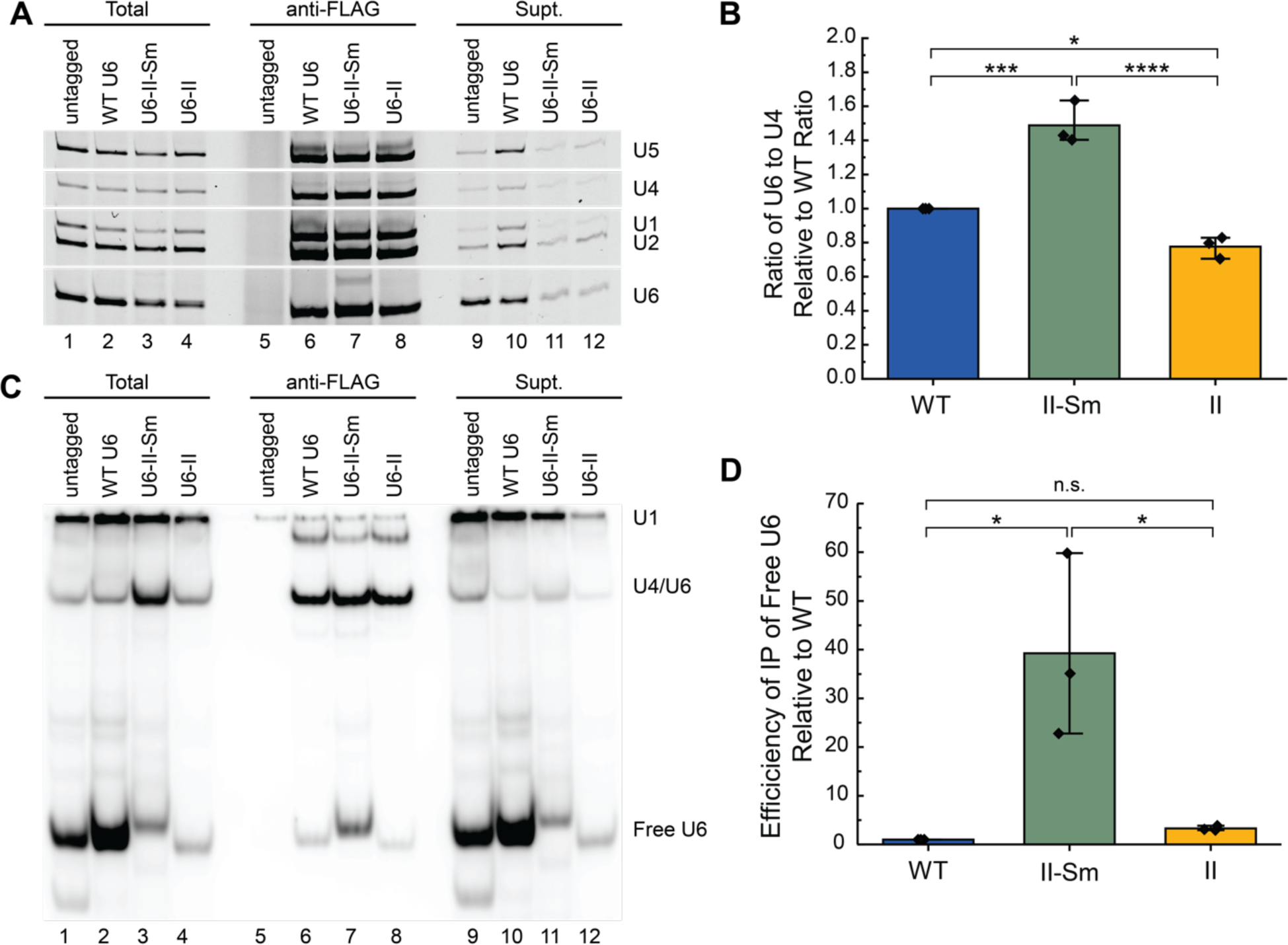
Free U6-II-Sm but not U6-II co-IPs with the SmD1 protein. **A)** Extracts from SmD1-3xFLAG strains or a control lacking the 3xFLAG epitope tag were immunoprecipitated using anti-FLAG antibodies. RNAs from the immunoprecipitate, supernatant, or total extract were extracted using phenol, precipitated, and analyzed by primer extension. **B)** Averages of the ratios of U6 to U4 in anti-FLAG IPs relative to WT samples from the experiment in Panel A (*n* = 3). **C)** Analysis of U6 base pairing interactions with U4 snRNA. RNAs from the immunoprecipitate, supernatant, or total extract were isolated under conditions which maintain intact base pairing. RNAs were then analyzed via hybridization with radiolabeled DNA probes complementary to U6 and U1 snRNAs. **D)** Average efficiencies of free U6 IP relative to WT U6 and after normalization to U1 from the experiment in Panel C (*n* = 3).

To confirm that increased co-IP of U6-II-Sm with Smd1 is due to Sm protein binding directly to the snRNA, we isolated RNAs from the co-IP under conditions that maintain base-pairing interactions to distinguish between U6 snRNAs co-IP’d as part of U4/U6 di-snRNA vs. free U6. U1, U4/U6, and U6 were detected and quantified by solution hybridization and native gel electrophoresis. As expected, IP’s from all three extracts contained base-paired U4/U6 (**Fig. 7C**, lanes 6-8). We quantified the efficiency of free U6 snRNP IP (anti-FLAG/Total) relative to wild type after normalizing to U1 snRNA because the amount of free U6 snRNA varied greatly between the extracts (**Fig. 7C**, lanes 2-4). Anti-FLAG-Smd1 samples from strains expressing U6-II-Sm showed a ∼40-fold higher efficiency of free U6 (U6 snRNP) IP compared to WT while those from strains expressing U6-II showed no significant increase over WT (**Fig. 7C, D**). Furthermore, the average ratio of U4/U6 to free U6 in the U6-II-Sm Smd1 IP sample is 1.8 (**Fig. 7C**, lane 7), close to the 2:1 ratio predicted from **Fig. 7B**. Thus, we conclude that the strong stabilization of U6-II by addition of an Sm-binding site is due to direct binding of the Sm ring.

## DISCUSSION

Eukaryotic RNAPs I, II, and III are specialist enzymes that transcribe specific sets of genes, each with defined characteristics and unique downstream post-transcriptional processing steps. Few examples exist within the literature of genes that are actively transcribed by different polymerases in different organisms or have been manipulated to switch transcription from one polymerase to another (Gunnery and Mathews 1995; Dergai and Hernandez 2019). Herein, we have shown that an snRNA central to pre-mRNA splicing, U6, is unexpectedly adaptable in both the route of its biosynthesis and its cellular interaction partners. We have shown that functional U6 can be synthesized by RNAP II under direction of the U4 snRNA promoter and terminator sequences in *S. cerevisiae*. However, high instability of the U6-II transcript was observed. Addition of an Sm binding site to U6-II increased stability, likely as the result of Sm ring binding. Uncoupling U6 synthesis from RNAP III allows for investigation of processing steps solely dependent on recognition of elements within the U6 snRNA rather than dependence on RNAP III transcription and provides the ability to control U6 snRNA synthesis via regulated RNAP II promoters.

### Formation of a U6•Sm snRNP

The U6-II-Sm snRNA is bound by both Lsm and Sm proteins but we do not know if it can be bound by both the Lsm and Sm complexes simultaneously, if one complex is exchanged for the other during U6 biogenesis or splicing, or if one complex is more likely to be involved in catalytically-competent spliceosome formation over the other. Nonetheless, this observation indicates flexibility in what proteins interact with the 3’ region of U6.

Sequential exchange of Sm for Lsm protein complexes on an RNA has previously been observed for the *S. pombe* telomerase RNA, TER1 (Tang et al. 2012). In this case, TER1 RNA initially is bound by Sm proteins, which facilitate its cleavage by the spliceosome and hypermethylation of its RNA cap. The Sm ring is then replaced by the Lsm ring for assembly of TER1 into telomerase RNPs. In the case of U6-II-Sm in *S. cerevisiae*, it is possible that the Sm ring provides 3’ end protection during nuclear-cytoplasmic shuttling (Becker et al. 2021). Whether or not the Sm and Lsm complexes need to be exchanged to make functional spliceosomes is not yet clear nor have we excluded the possibility of simultaneous occupancy of U6 by both complexes. Inspection of the yeast U6 snRNP structure (Montemayor et al. 2018) suggests it is possible that the Sm and Lsm rings can bind U6-II-Sm simultaneously as the Sm binding site should not be occluded by the Lsm ring or U6 snRNA secondary structure.

The Lsm ring on WT U6 is released during formation of the spliceosome active site to allow binding of the NineTeen Complex (NTC) (Townsend et al. 2020). If Sm or Lsm complexes are retained on U6-II-Sm snRNAs during splicing, it is likely that this can only be accomplished by major remodeling of essential splicing factors and formation of spliceosome architectures distinct from those currently known. For splicing to proceed normally, we would therefore expect that the Sm ring on U6-II-Sm would also be released during spliceosome activation. The Sm rings on U1, U2, U4, and U5 snRNAs are not released at any point during splicing nor is it clear how the Sm•snRNA interaction would be disassembled. It is therefore intriguing to speculate that if U6-II-Sm RNPs are incorporated into spliceosomes, the same mechanism responsible for Lsm release can also dissociate a U6•Sm complex (or both U6•Sm and U6•Lsm interactions if both rings are simultaneously present). Inefficient dissociation of the U6•Sm interaction during activation may be the origin of the reduced splicing efficiency we observed for the U3 pre-mRNAs. Since the Lsm2-8 ring also facilitates U4/U6 di-snRNP formation (Didychuk et al. 2015), it is possible that defects in this step also contribute to the reduced splicing, but this seems less likely since solution hybridization assays indicate enhanced levels of U4/U6 di-snRNAs in U6-II-Sm extracts (**Fig. 7C**).

### Conclusions

It remains to be determined what evolutionary advantage, if any, is conferred by RNAP III specialization of U6 snRNA synthesis over RNAP II. It is possible that transcription by RNAP III is a relic of the mechanism by which U6 snRNA was originally acquired by the eukaryotic genome from Group II self-splicing introns (Novikova and Belfort 2017). A relevant parallel is that all yeast small nucleolar RNAs are synthesized by RNAP II except snR52, which is synthesized by RNAP III (Harismendy et al. 2003; Moqtaderi and Struhl 2004). Integration of transcription-factor like subunits into the RNAP III core during evolution may have promoted more efficient and rapid transcription, facilitating production of the highly abundant ncRNAs synthesized by RNAP III (Barba-Aliaga et al. 2021).

Regardless of the unanswered evolutionary questions, our system for synthesis of U6 by RNAP II in yeast provides a useful tool for dissecting how *cis*-acting sequences within U6 shape its biogenesis without the transcriptional restraints imposed by RNAP III regulatory elements found within the *SNR6* gene. For example, U6 synthesis independent of the A block internal promoter element will enable sequence manipulation of the 5’ stem-loop *in vivo*, and rapid induction or repression of U6-II synthesis using the galactose inducible promoter will allow conditional expression of mutant U6 alleles. As the 5’ stem-loop is the sole region of U6 with no known function in spliceosome function, it is a promising target for engineering affinity tags or binding sites for fluorophores or fluorescent proteins.

## SUPPLEMENTAL MATERIAL

Supplemental material is available for this article.

## ACKNOWLEDGEMENTS

We thank Beate Schwer for providing plasmids for expression of Lsm5 and Lsm8 and Reinhard Lührmann for a gift of H20 monoclonal antibody. We thank Allison Didychuk for critical comments on the manuscript and members of the Brow, Butcher, and Hoskins labs for helpful discussions.

## FUNDING

This work was supported by grants from the National Institutes of Health (R35 GM136261 to AAH and R35 GM118075 to DAB).

## COMPETING INTERESTS

AAH is a member of the scientific advisory board and carrying out sponsored research for Remix Therapeutics.

## MATERIALS AND METHODS

### Yeast Strains and Growth

Yeast strains used in these experiments are described in **Supplemental Table 1**. Yeast were grown in standard media as indicated in the figure legends (Treco and Lundblad 1993). Plasmid transformations and shuffling assays were carried out using standard procedures as described (Sikorski and Boeke 1991).

Strains carrying either a C-terminal 2xV5 epitope tag on Lsm8 or a C-terminal 3xFLAG epitope tag on Smd1 were created using PCR-based tagging (Janke et al. 2004). Briefly, synthetic, linear DNA fragments coding for the 2xV5 or 3xFLAG epitope tags (IDT) were digested with HindIII and SalI and then ligated into those same restriction sites of a digested pAG32 plasmid, which also contained an expression cassette for the yeast hygromycin resistance (HygR) marker (Goldstein and McCusker 1999). The resultant plasmids were then used as templates for PCR to prepare linear DNAs for gene tagging by homologous recombination. After transformation of the DNA fragments, yeast containing the integration cassettes were selected by growth on YPD plates hygromycin (300 µg/mL; Sigma Aldrich). Colonies were screened by PCR to confirm proper integration of the tag at the correct chromosomal location.

### Oligonucleotides

The sequences for the DNA oligos used for cloning, primer extension, Northern blotting, and solution hybridization assays are shown in **Supplementary Table 2**.

### Plasmids

Plasmids used in these experiments are listed in **Supplemental Table 3**. To make pRS314-SNR14-6-14, the *SNR14* promoter from pRS313-SNR14 (Kuehner and Brow 2006) was amplified by PCR with primers PsnR14(223)-Xho-F and U4-U6-R, and the XhoI/NruI-digested product was ligated to XhoI/NruI-digested pRS314-554H6 (Martin et al. 2001) to create pRS314-SNR14-6. Then, recombinant PCR (Horton et al. 1993) was used to replace the *SNR6* downstream DNA with *SNR14* downstream DNA. The first round of PCR used primers PsnR14(223)-Xho-F and U4(12)-U6(24)-R on pRS314-SNR14-6, and primers U6(12)-U4(23)F and snR14-BamHI-701R on pRS313-SNR14. These products were then joined and amplified by PCR with PsnR14(223)-Xho-F and snR14-BamHI-701R, and the digested product was ligated into XhoI/BamHI-cut pRS314.

pRS314-SNR6-II was created by recombinant PCR as described above but using primer pairs PsnR14(223)-Xho-F/Rnt1(12)-U6(24)R and U6(12)-Rnt1(23)F/snR14-BamHI-701R in the first round with pRS314-SNR14-6-14 as template.

pRS314-GAL-SNR6-II was created by recombinant PCR. The first round of PCR used primers GAL1-XhoI-608F and U4(14)-GAL1(24)R to amplify the *GAL1* upstream region from genomic DNA prepared from yeast strain PJ43-2b (Kaiser and Brow 1995), and primers Gal(11)-U4(25)F and snR14-KpnI-701R to amplify pRS424-SNR6-II. These products were then joined and amplified by PCR with primers GAL1-XhoI-608F and snR14-KpnI-701R, and the digested product was ligated into XhoI/KpnI-cut pRS314.

pRS314-GAL-SNR6-II-Sm was created by recombinant PCR using the following primer pairs and pRS314-GAL-SNR6-II as template: GAL1-XhoI-608F/Sew-U6::U4R and Sew-U6::U4-F/snR14-KpnI-701R.

High copy versions of these plasmids were made by subcloning the inserts into pRS424.

### Northern blotting and primer extension

Total cellular RNA was isolated by phenol extraction in the presence of guanidinium thiocyanate (Wise et al. 1983). Northern blot analysis used 25 µg of RNA per lane of a 20 cm x 20 cm x 1.5 mm 6% polyacrylamide, 8.3 M urea gel run in 50 mM TBE buffer at 400 V for 1.5 h. The gel was washed 2 x 15 min in Transfer buffer (12 mM Tris acetate, pH 7.6, 0.3 mM EDTA) at 4°C and transferred overnight to a Zeta-Probe nylon membrane (Bio-Rad) at 20 V and 4°C in a Hoefer Transphor TE 42 unit. The blot was hybridized to a ^32^P-labeled oligonucleotide complementary to U6 RNA (U6D, Table S3) in Rapid-hyb buffer (Amersham) according to the manufacturer’s instructions.

Primer extension analysis used 4 µg total cellular RNA per reaction combined with 200 fmol of each 5’-^32^P-labeled oligonucleotide (U1-SH, U4-14B, U3 21-mer, U6D or U6B) in a total volume of 2.5 µL of Annealing Buffer (10 mM TrisCl, pH 8.0, 250 mM KCl). Mixtures were incubated at 90°C for 3 min, on ice for 3 min, and at 45°C for 5 min. 6.5 µL of RT mix (35 mM TrisCl, pH 8.0, 11.5 mM MgCl_2_, 11.5 mM DTT, 0.6 mM each dNTP, 4.5 U AMV reverse transcriptase) was added and the reaction incubated at 45°C for 15 min. 2.5 µL of formamide loading buffer was added and the mixture heated at 90°C for 3 min before loading on a 6% polyacrylamide (19:1) gel with 8.3 M urea and 50 mM TBE.

### TMG cap antibody gel shift

Ten µg of total cellular RNA per lane were analyzed as described previously (Kwan et al. 2000) using ^32^P-labeled oligonucleotides U6D or U4-14B.

### Nondenaturing RNA isolation

All steps were performed on ice with cold solutions. Ten ODs of log phase culture (at 0.6-0.8 OD_600_) was added to cold MilliQ water up to 50 mL. Cells were pelleted and washed with RNA Buffer (200 mM TrisCl pH 7.5, 100 mM EDTA, and 500 mM NaCl) before resuspension in 300 µL RNA buffer. 200 µL 425-600 micron acid-washed glass beads (Scientific Industries) and 300 µL 1:1 phenol/CHCl_3_ were added and cells were lysed by vortexing twice at high speed for 1 min with a 1 min rest on ice between. The aqueous phase was extracted again with an equal volume of 1:3 phenol/CHCl_3_ and precipitated with three volumes 100% ethanol at -20°C. RNA was pelleted and washed with 70% v/v ethanol before resuspension in RNase-free water.

### Solution hybridization

DNA probes (U1-SH, U4-14B, and U6-SH) were end labeled with [γ-^32^P]-ATP (PerkinElmer) using T4 PNK (Thermo Scientific) for 1 h at 37°C followed by enzyme inactivation by heating to 60°C for 20 min according to the manufacturer’s instructions. Probes were hybridized to 5 μg total RNA at 37°C for 15 min in Solution Hybridization Buffer (50 mM Tris pH 7.5, 1 mM EDTA, and 150 mM NaCl) before mixing 1:1 with 50% v/v cold glycerol (22 µL total volume). Twenty µL of the reaction was loaded onto a 20 cm x 22 cm x 1 mm 9% nondenaturing polyacrylamide gel in 50 mM TBE. RNA:DNA hybrids were separated for 5 hours at 300V at 4°C. The gel was dried under vacuum for 30 min before exposure to a phosphor screen overnight followed by imaging (Li and Brow 1993).

### Measurement of cell growth and U6 RNA stability under repressing conditions

Yeast cells were grown at 30°C in YEP with 2% galactose to OD600 = 0.5 – 1.0, pelleted, washed with sterile water, and resuspended to the same density in YEP with 2% glucose. Growth curves were generated by continually diluting cultures as required to keep the OD_600_ between 0.2 and 1.0 (Patterson and Guthrie 1987). Aliquots (50 mL) of the culture were collected at the indicated times to prepare RNA for primer extension.

### Analysis of snRNPs by glycerol gradients

Yeast whole cell extracts (yWCE) were prepared using the liquid N2 method and processing with a ball mill (Mayas et al. 2006). yWCEs in dialysis buffer (25 mM HEPES-KOH pH 7.9, 50 mM KCl, 20% v/v glycerol, 0.5 mM DTT) were adjusted to 8% v/v glycerol and 500 µL of the dilution was layered onto 12.5 mL 10-30% (v/v) linear glycerol gradients (Gradient Master) in 20 mM HEPES pH 7.9, 150 mM KCl, 1 mM Mg(OAc)_2_, 0.1% v/v NP40, 0.5 mM PMSF, 1 µg/ml Pepstatin A, and 1 µg/ml Leupeptin. Sedimentation was performed in a Beckman SW40 rotor for 22 h at 35,000 rpm, 4°C. Fractions of 500 µL were collected from the top.

RNA was isolated from glycerol gradient fractions with a 25:24:1 phenol/CHCl_3_/isoamyl alcohol extraction and precipitated for 30 min at -20°C with 0.1 volume 3M NaOAc, 10 µg tRNA, and 2.5 volumes 100% EtOH. Precipitated RNA pellets were washed with 96% EtOH and resuspended in 8 µL dH_2_O. Total RNA samples were prepared from 50 µL yWCE resuspended in an equal amount of Splicing Dilution Buffer (100 mM TrisCl pH 7.5, 10 mM EDTA pH 8.0, 1% SDS (w/v), 150 mM NaCl, 0.3 M NaOAc pH 5.2-5.4) and 100 µL dH_2_O. After incubation at 37°C for 15 min, 200uL dH_2_O was added to samples. Total RNA was extracted with 400 uL 25:24:1 phenol/CHCl_3_/isoamyl alcohol four times and precipitated as described above. Pellets were resuspended in 32 µL dH_2_O. Primer extension of isolated RNA was accomplished with the qScript Flex cDNA Synthesis Kit using 8 µL volumes with 5’ labeled Dy682 primers (U1RT136, U2RTALL124, U4RTALL, U5, U6D). Amplified cDNA products were separated on a 7% denaturing PAGE gel for 1 hr at 22 W and imaged on an Amersham Typhoon NIR laser scanner (Cytiva).

### Immunoprecipitation

yWCE for IP were prepared using the liquid N2 method and processing with a ball mill (Mayas et al. 2006) without dialysis. A 20 µL volume of protein G magnetic beads (BioRad SureBeads™ #1614023) per IP reaction was washed three times with 1.5 mL cold IPP150 (150 mM NaCl, 0.1% v/v NP-40 Surfact-Amps™ Detergent Solution, 10 mM Tris pH 8.0, and 0.1% w/v azide) and three times with 1.25 mL cold IPP500 (500 mM NaCl, 0.1% NP-40 Surfact-Amps™ Detergent Solution, 10 mM Tris pH 8.0, and 0.1% w/v azide). Beads were coupled with 4.8 µg of either anti-V5 (ThermoFisher Scientific, R960-25) or anti-FLAG (Millipore Sigma, F1804) antibody overnight at 4°C in 100 µL total volume of magnetic bead suspension in IPP150.

After incubation, beads were washed three times with 1.5 mL of IPP150 and resuspended in 80 µL IPP150. IP reactions contained the magnetic bead suspension, 40 µL yWCE, 1.25 µL RNasin® Plus Ribonuclease Inhibitor (Promega, #N2611), 2.5 mM DTT, and IPP150 to 200 µL total volume. Samples of total RNA were assembled without beads and incubated as described. Reactions were agitated overnight at 4°C. Supernatant from reactions was saved and beads were washed five times in IPP150 containing 2.5 mM DTT. Protein bound RNA was released by Proteinase K incubation (0.8 µg/µL Proteinase K (Thermo Scientific), 50 mM Tris pH 7.5, 10 mM EDTA, 1% w/v SDS, and RNase-free water to 200 µL) at 37°C for 30 minutes. The supernatant was transferred to a fresh tube and 200 µL of IPP150 with 0.1 mg/mL *E. coli* tRNA (Roche) was added. To the IP supernatant and total RNA samples, 200 µL IPP150 containing 10 mM EDTA, 0.5% w/v SDS, and 0.1 mg/mL *E. coli* tRNA was added.

RNA was extracted once with phenol:chloroform and precipitated with 300 µL 3M NaOAc pH 5.2 and 1 mL ice cold 100% EtOH overnight at -80°C. RNA was pelleted for 30 min at 14K, 4°C and washed with 70% EtOH. Dried pellets were resuspended in 10-12 µL volumes of RNase free water. Immunoprecipitated RNAs were reverse transcribed into cDNA using SuperScript™ III Reverse Transcriptase (ThermoFischer Scientific, # 18080044) according to the manufacturer’s instructions using 0.2 pmol of each primer (U1RT136, U2RTALL124, U4RTALL, U5, U6B labeled with 5’ IRDye® 700, IDT; see Table S2). Reactions were diluted in one volume of clear formamide loading dye and heat denatured at 95°C before loading onto a 22 cm x 22 cm x 0.75 mm denaturing 7% 19:1 PAGE gel. Fluorescent cDNA products were separated for 80 min at 35 W and imaged on an Amersham Typhoon NIR laser scanner (Cytiva). U6 to U4 ratios of the IP based on primer extension analysis (Fig. 7A) and reported in Fig. 7B were calculated using Equation 1. The *IP* subscript represents band intensities determined from the anti-FLAG samples.

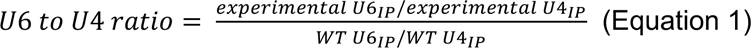

For RNAs isolated by IP and analyzed by solution hybridization, RNAs were isolated as described in the preceding paragraph, but with no NaOAc added. Efficiency of free U6 IP relative to wild type was calculated using Equation 2 after normalization of the samples using the U1 snRNA bands.

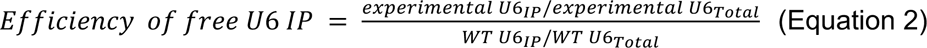

### Statistical Analysis

All statistical tests were conducted using R (R Core Team, 2021). Data collected from solution hybridization experiments was assumed to be normally distributed. One-way ANOVA was applied to test for differences in sample means between the WT, U6-II, and U6-II-Sm experimental strains in solution hybridization and co-IP experiments. Means between two experimental groups within the dataset were compared with a post-hoc Tukey multiple pairwise-comparisons test to determine the statistical significance level (p = 0.05*, 0.01**, 0.001***, 0.0001****).

