## Supplemental Figures and Tables for "Yeast U6 snRNA made by RNA polymerase II is less stable but functional"

**CORRESPONDING AUTHORS:**

**AFILLIATIONS:**

[1]Department of Chemistry, University of Wisconsin-Madison, Madison, WI 53706

[2]Department of Biomolecular Chemistry, University of Wisconsin School of Medicine and Public Health, Madison, WI 53706

[3]Department of Biochemistry, University of Wisconsin-Madison, Madison, WI 53706

**CONTENTS:** 4 Supplemental Figures
3 Supplemental Tables
Supplemental References

### Supplemental Figures

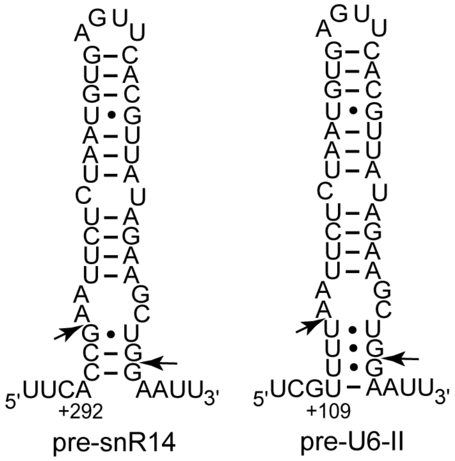


**Figure S1.** Predicted positions of Rnt1 cleavage sites (arrows) in the primary transcript of SNR6-II (right) compared to the previously mapped cleavage sites in pre-U4 snRNA (left; Allmang et al. 1999). The first 135 base pairs downstream of U6 position +112 were deleted from the SNR14-6-14 allele to align the mature 3' end of the shortest version of yeast U6 snRNA with the upstream Rnt1 cleavage site (see Figure 1B).

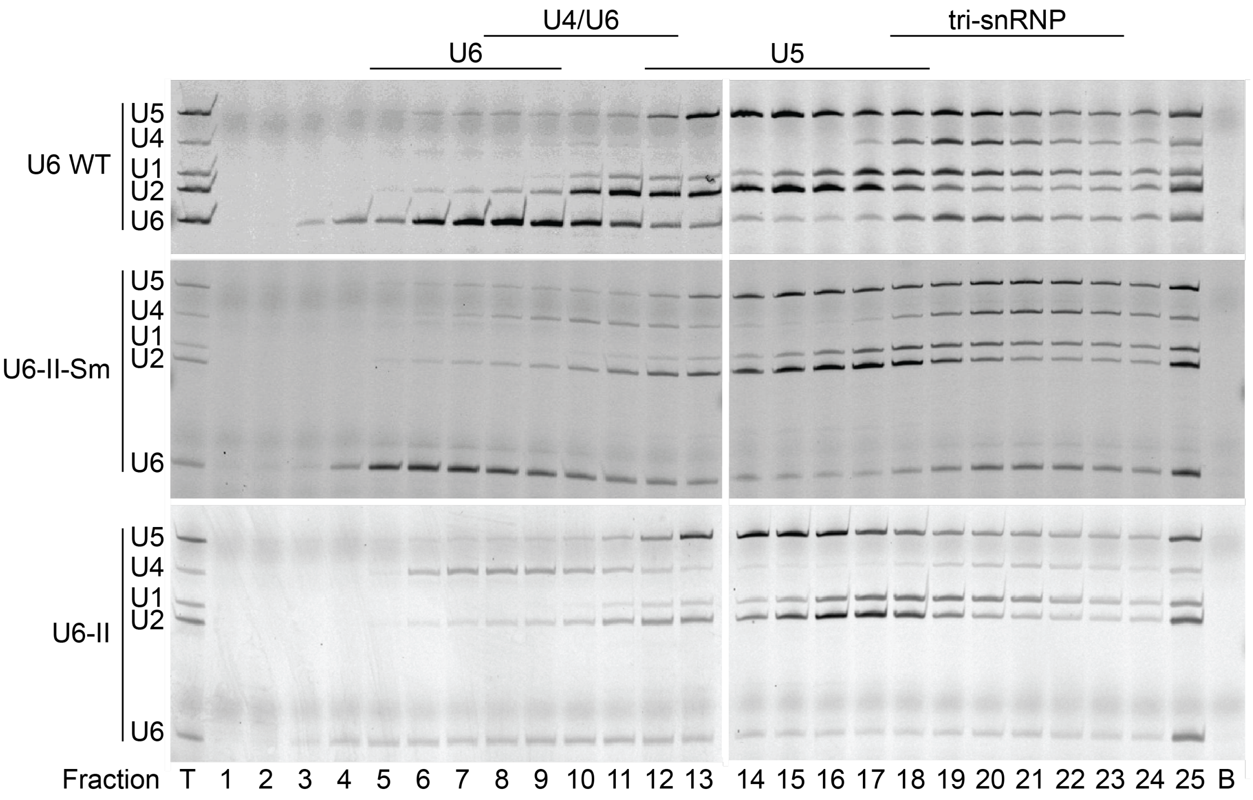


**Figure S2. Primer extension of snRNAs from glycerol gradient fractions.** RNAs were isolated from fractions taken from glycerol gradient sedimentation. Levels of primer extension products were quantitated as seen in Figure 5.

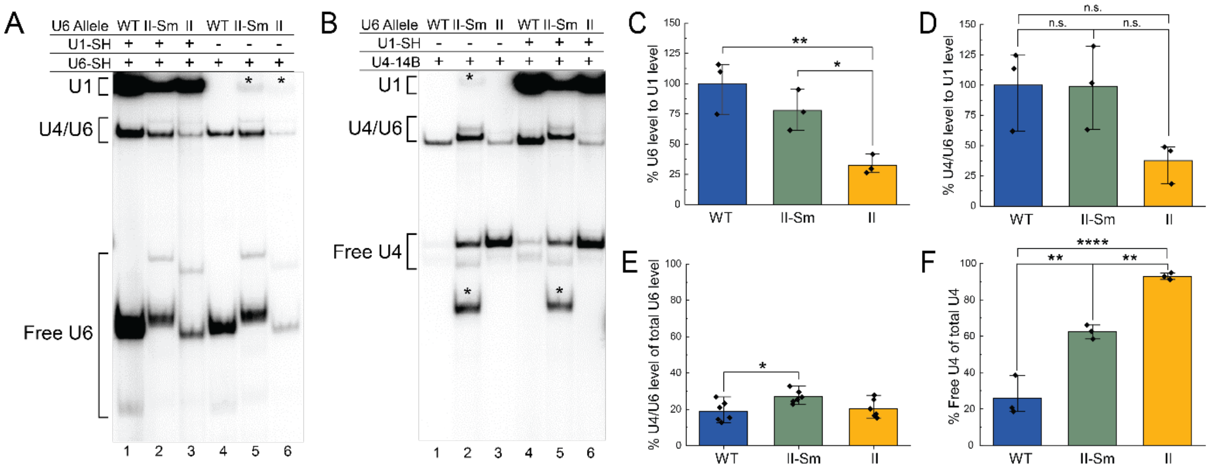


**Figure S3. Free U4 snRNA accumulates and U4/U6 is diminished in yeast cells producing U6-II. A, B)** Non-denaturing gel electrophoresis of cold-extracted whole-cell RNA from strains bearing the indicated U6 allele hybridized to probes against the indicated snRNAs. Positions of U4/U6 di-snRNA and free U1, U4, and U6 snRNAs are indicated. The bands indicated with asterisks are believed to be U6-II-Sm snRNA hybridized with the U4-14B probe due to inclusion of the U4 Sm binding site. **C, D, E, F)** Quantification of di-snRNA ratios and snRNA abundance from gel bands present in solution hybridization assays. di-snRNA ratios and relative abundance were calculated from three biological replicates. Band volume for U1 was adjusted to exclude contributions from a faint band in U6-II-Sm lanes running at the same height as U1. Sample means were compared with one-way ANOVA followed by a post hoc Tukey multi-pairwise analysis. ($p$ = 0.05*, 0.01**, 0.001***, 0.0001****)

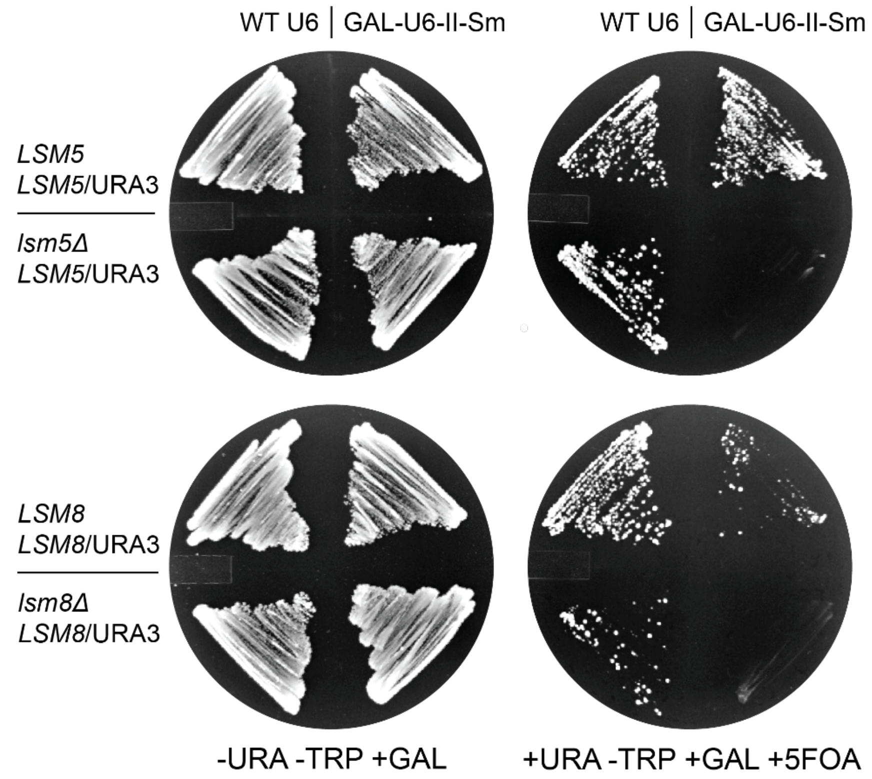


**Figure S4. Genetic interaction between U6-II-Sm and Lsm Proteins.** *lsm5* or *lsm8* genes were deleted from strains expressing either WT U6 or U6-II-Sm RNAs and complemented with Lsm5 or Lsm8 expression from a URA3/CEN-marked plasmid. These deletion strains or their corresponding controls were then grown on medium selecting for (-URA -TRP +GAL) or against (+URA -TRP +GAL +5FOA) the *URA3*-marked plasmid bearing the WT *LSM* gene. Deletion of the *lsm5* or *lsm8* genes and loss of the corresponding URA3-marked plasmid is lethal in the presence of U6-II-Sm but not WT U6.

**Supplementary Table 1: Yeast strains**

| Strain | Parent | Genotype | Reference |
|---|---|---|---|
| MWK027 | PJ43-2b | MATα ura3-52 trp1-1 his3-11,15 leu2-3,112 ade2-1 lys2Δ2 met2Δ1 can1-101 snr6::LEU2 [YCp50-ΨWT] | Kaiser & Brow 1995 |
| CJM000 | ANK640 | MATa, snr6::LEU2, snr14::trp1::ADE2, trp1, ura3, lys2, his3, ade2, [pRS316-U4wt-U6mini] | McManus et al. 2007 |
| DAB101 | CJM000 | MATa, snr6::LEU2, snr14::trp1::ADE2, trp1, ura3, lys2, his3, ade2, [pRS313-U4wt + pRS314-U6wt] | This work |
| yAAH1361 | CJM000 | MATa, snr6::LEU2, snr14::trp1::ADE2, trp1, ura3, lys2, his3, ade2, [pRS313-U4wt + pRS314-U6-II-Sm] | This work |
| yAAH1442 | CJM000 | MATa, snr6::LEU2, snr14::trp1::ADE2, trp1, ura3, lys2, his3, ade2, [pRS313-U4wt + pRS314-U6-II] | This work |
| yAAH2687 | BY4743 | MATa/α his3Δ1/his3Δ1 leu2Δ0/leu2Δ0 LYS2/lys2Δ0 met15Δ0/MET15 ura3Δ0/ura3Δ0 LSM5::kanMX [pRS316-LSM5] | Roth et al. 2018 |
| yAAH2690 | CJM000 | MATa, snr6::LEU2, snr14::trp1::ADE2, trp1, ura3, lys2, his3, ade2, SMD1::3xFLAG-HygR [pRS316-U4wt-U6mini] | This work |
| yAAH2719 | CJM000 | MATa, snr6::LEU2, snr14::trp1::ADE2, trp1, ura3, lys2, his3, ade2, LSM8::2xV5-HygR, [pRS316-U4wt-U6mini] | This work |
| yAAH2720 | CJM000 | MATa, snr6::LEU2, snr14::trp1::ADE2, trp1, ura3, lys2, his3, ade2, [pRS313-U4wt + pRS314-U6wt + pRS416-LSM5] | This work |
| yAAH2721 | CJM000 | MATa, snr6::LEU2, snr14::trp1::ADE2, trp1, ura3, lys2, his3, ade2, [pRS313-U4wt + pRS314-U6wt + pRS416-LSM8] | This work |

| | | | |
|---|---|---|---|
| yAAH2722 | CJM000 | MATa, snr6::LEU2, snr14::trp1::ADE2, trp1, ura3, lys2, his3, ade2, [pRS313-U4wt + pRS314-U6-II-Sm + pRS416-LSM5] | This work |
| yAAH2723 | CJM000 | MATa, snr6::LEU2, snr14::trp1::ADE2, trp1, ura3, lys2, his3, ade2, [pRS313-U4wt + pRS314-U6-II-Sm + pRS416-LSM8] | This work |
| yAAH2736 | CJM000 | MATa, snr6::LEU2, snr14::trp1::ADE2, trp1, ura3, lys2, his3, ade2, LSM5Δ::KanMX [pRS313-U4wt + pRS314-U6wt + pRS416-LSM5] | This work |
| yAAH2737 | CJM000 | MATa, snr6::LEU2, snr14::trp1::ADE2, trp1, ura3, lys2, his3, ade2, LSM8Δ:: HygR [pRS313-U4wt + pRS314-U6wt + pRS416-LSM8] | This work |
| yAAH2750 | CJM000 | MATa, snr6::LEU2, snr14::trp1::ADE2, trp1, ura3, lys2, his3, ade2, LSM5Δ::KanMX [pRS313-U4wt + pRS314-U6-II-Sm + pRS416-LSM5] | This work |
| yAAH2751 | CJM000 | MATa, snr6::LEU2, snr14::trp1::ADE2, trp1, ura3, lys2, his3, ade2, LSM5Δ:: HygR [pRS313-U4wt + pRS314-U6-II-Sm + pRS416-LSM5] | This work |
| yAAH2773 | CJM000 | MATa, snr6::LEU2, snr14::trp1::ADE2, trp1, ura3, lys2, his3, ade2, LSM8::2xV5 HygR, [pRS313-U4wt + pRS314-U6WT] | This work |
| yAAH2774 | CJM000 | MATa, snr6::LEU2, snr14::trp1::ADE2, trp1, ura3, lys2, his3, ade2, LSM8::2xV5 HygR, [pRS313-U4wt + pRS314-U6-II-Sm] | This work |
| yAAH2775 | CJM000 | MATa, snr6::LEU2, snr14::trp1::ADE2, trp1, ura3, lys2, his3, ade2, LSM8::2xV5 HygR, [pRS313-U4wt + pRS314-U6-II] | This work |
| yAAH2853 | CJM000 | MATa, snr6::LEU2, snr14::trp1::ADE2, trp1, ura3, lys2, his3, ade2, SMD1::3xFLAG HygR, [pRS313-U4wt + pRS314-U6wt] | This work |

| | | | |
|---|---|---|---|
| yAAH2854 | CJM000 | MATa, snr6::LEU2, snr14::trp1::ADE2, trp1, ura3, lys2, his3, ade2, SMD1::3xFLAG HygR, [pRS313-U4wt + pRS314-U6-II-Sm] | This work |
| yAAH2855 | CJM000 | MATa, snr6::LEU2, snr14::trp1::ADE2, trp1, ura3, lys2, his3, ade2, SMD1::3xFLAG HygR, [pRS313-U4wt + pRS314-U6-II] | This work |

### Supplementary Table 2: Oligonucleotides

| Oligo | Sequence (5' to 3') |
|---|---|
| PsnR14(223)-Xho-F | CCGCTCGAGTAAGTAACCTCTGCATTGTC |
| U4-U6-R | CCCTCGCGAACGGAGTATTAAGGAAGGAAGTG |
| U4(12)-U6(24)-R | TCGGTAATGAAAAAAACGAAATAAATCTCTTTGTAA |
| U6(12)-U4(23)F | TTATTTCGTTTTTTTCATTACCGATATTCATTCTT |
| snR14-BamHI-701R | CGGGATCCTTCCTCTCTGCTGTTTTAG |
| Rnt1(12)-U6(24)R | CATTAGAGAATTAAAACGAAATAAATCTCTTTGTAA |
| U6(12)-Rnt(23)F | TTATTTCGTTTTAATTCTCTAATGTGAGTTCACGT |
| GAL1-XhoI-608F | CCGCTCGAGATCATATTACATGGCATTACCA |
| U4(14)-GAL1(24)R | CTGTTCCTTTTATATCTGTTAATAGATCAAAAATCATC |
| GAL1(11)-U4(25)F | CTATTAACAGATATAAAAGGAACAGAATATTAGTTA |
| snR14-Kpn1-701R | CGGGGTACCTTCCTCTCTGCTGTTTTAG |
| Sew-U6::U4-R | AAAAGGTATTCCAAAAATTCTTTGTAAAACGGTTCATCCTT |
| Sew-U6::U4-F | CAAAGAATTTTTGGAATACCTTTTAATTCTCTAATGTGAGTTCA |
| U1-SH | CCGTATGTGTGTGTGACC |
| U4-14B | AGGTATTCCAAAAATTCCC |
| U6-SH | ATTGTTTCAAATTGACCAAAT |
| U1RT136 | GAC CAA GGA GTT TGC ATC AAT GAC |
| U2RTALL124 | TTT GGG TGC CAA AAA ATG TGT ATT GTA |
| U4RTALL | GGT ATT CCA AAA ATT CCC TAC ATA GTC |
| U5 | AAG TTC CAA AAA ATA TGG CAA GC |
| U6B | TCATCCTTATGCAGGG |
| U6D | AAA ACG AAA TAA ATC TCT TTG |
| U3 21-mer | CCAAGTTGGATTCAGTGGCTC |

57 **Supplementary Table 3: Plasmids**

| Plasmid | Description | Reference |
|---|---|---|
| pRS314 | *TRP1*-marked, low-copy yeast shuttle plasmid | Sikorski and Hieter 1989 |
| pRS424 | *TRP1*-marked, high-copy yeast shuttle plasmid | Christianson et al. 1992 |
| pRS313-SNR14 | *SNR14* -224 to +701 cloned into the BamHI site | Kuehner and Brow 2006 |
| pRS314-14-6-14 | See Materials and Methods | This work |
| pRS424-14-6-14 | See Materials and Methods | This work |
| pRS314-SNR6-II | See Materials and Methods | This work |
| pRS424-SNR6-II | See Materials and Methods | This work |
| pRS314-GAL-SNR6-II | See Materials and Methods | This work |
| pRS314-GAL-SNR6-II-Sm | See Materials and Methods | This work |
| pAAH0229 | pRS314-GAL-SNR6-SM (U6-II-Sm)/TRP/AMP | This work |
| pAAH0412 | pRS314 WT U6/TRP/AMP | McManus et al. 2007 |
| pAAH0413 | pRS313 WT U4/HIS/AMP | McManus et al. 2007 |
| pAAH0667 | pRS314-GAL-SNR6-II (U6-II)/TRP/AMP | This work |
| pAAH1285 | pRS416 LSM5/URA/CEN | Roth et al. 2018 |
| pAAH1286 | pRS416 LSM8/URA/CEN | Roth et al. 2018 |

58

59
